# B cells sustain tumor-specific CD4 T cells to promote response to PD-1 targeted therapy

**DOI:** 10.64898/2026.09.18.751292

**Authors:** Abishek Vaidya, Etienne Humblin, Nataliya Prokhnevska, Verena van der Heide, Laurine Binet, Jachym Harwood, Raphaël Mattiuz, Anthony Lozano, Pauline Hamon, Bruno Cogliati, Simon Goldstein, Isabel Korpas, Katherine E. Lindblad, Suhaana Sriram, Harald Hartweger, Glaucia Furtado, Sacha Gnjatic, Edgar Gonzalez-Kozlova, Brad R. Rosenberg, Amaia Lujambio, Miriam Merad, Michel C. Nussenzweig, Alice O. Kamphorst

## Abstract

Immunotherapy transformed cancer treatment, yet precise correlates of response remain to be defined. While intratumoral follicular helper like (Tfhl) CD4 T cells and IgG1^+^ B cells have been associated with improved outcomes to PD-1 blockade for hepatocellular carcinoma (HCC), their mechanisms contributing to response are unclear. To address this question, we developed a murine model of HCC and examined CD8, CD4 and B cell responses. Increasing CD4-help sensitizes mice to PD-1 blockade, in a CD4- and CD8-dependent manner. Tumor-specific CD4 T cells contained Tfhl and Th1 populations, and both *Bcl6* and T-bet were required for efficacy. Antigen-specific B cells were essential for CD4-helper expansion, yet secretion of antibodies was not required for long-term survival. Thus, B cells are critical for effective immunotherapy, but secreted antibodies are not.

## MAIN TEXT

Immune checkpoint blockade (ICB), particularly PD-1/PD-L1 inhibition, has transformed the treatment landscape of several solid tumors, producing durable responses in melanoma, non-small cell lung cancer, renal cell carcinoma and other malignancies (*1, 2*). Although CD8 T cells are widely regarded as the principal mediators of tumor regression (*3, 4*), many tumors harbor abundant CD8 T cells yet fail to respond to ICB, indicating that tumor control also depends on other immune cells, particularly CD4-helper T cells (*5–8*). Enhancement of tumor-specific CD8 T cell responses by CD4-help has been reported in several animal models (*9–15*). However, most of these studies have focused on T cell priming and/or T cell responses induced by vaccination strategies.

CD4 T cells orchestrate multiple layers of adaptive immunity: CD4 helpers activate dendritic cells (DCs) through CD40 ligation improving productive CD8 T cell priming (*16–20*); produce cytokines (such as IL-2 and IL-21) critical for CD8 differentiation and survival (*13, 14, 21, 22*); promote B cell activation, isotype switching and selection of high affinity B cells (through CD40-CD40L axis and cytokines)(*23*). In murine tumors, CD4 T helper type 1 cells (Th1) which secrete IFN-γ and express the transcription factor T-bet are more often associated with effective anti-tumor activity (*9, 24, 25*). However, in patients, CD4 T cells recognizing tumor antigens are characterized by transcriptional programs associated with chronic antigen stimulation and similarities to T follicular helper cells (Tfh) which aid B cells (*26, 27*).

In hepatocellular carcinoma (HCC) patients receiving neoadjuvant PD-1 blockade, we reported that tumor necrosis was associated with intratumoral expansion of Tfh-like (Tfhl) CD4 T cells and PD-1^hi^ CD8 T cells with effector features (*8*). Spatial analysis revealed triads of Tfhl, activated DCs and progenitor-exhausted PD-1^+^ CD8 T cells (TXp) in responder patients, suggesting that interactions between Tfhl and DCs may enhance differentiation of TXp into effector-like PD-1^hi^ CD8 T cells to enable tumor control. Tfhl frequency also correlated with plasma cell abundance (*8*), and recent analysis uncovered marked expansion of intratumoral IgG1^+^ plasma cells in HCC patients responding to PD-1 blockade (*28*). Enrichment of IgG1^+^ plasma cell signature also associated with response to ICB in melanoma and additional HCC cohorts (*28*). In 2020, three concurrent reports highlighted intratumoral B cells and tertiary lymphoid structures (TLS) as features of ICB response in several solid tumors (*29–31*). In soft-tissue sarcoma, B cells emerged as the strongest predictor of response, even after accounting for CD8 T cell density (*29*). These clinical associations underscore the need for a deeper understanding of the role of B cells in anti-tumor immunity.

To address these questions, we developed a murine model of HCC where we could track CD8, CD4 and B cell responses. Despite immune infiltration, this model recapitulates resistance to PD-1 blockade. However, enhancing CD4 T cell help rendered tumors responsive to PD-1 blockade. ICB-mediated survival was dependent on CD8 and CD4T cells, as well as B cells. Tumor-specific CD4 T cells differentiated into Th1 and Tfhl states and genetic perturbations revealed that both T-bet and *Bcl6* are required for CD4-help-mediated responses to ICB. Furthermore, although antigen-specific B cells were essential for tumor control, antibody secretion was dispensable. Our findings uncover a critical role of B cells but not their secreted antibody products in sustaining CD4-help and enabling effective responses to immune checkpoint blockade.

## RESULTS

### Autochthonous murine model of liver cancer with tractable CD8, CD4 and B cell responses

We modified a previously reported autochthonous HCC mouse model induced by hydrodynamic tail vein injection (HDTVI) of oncogenic plasmids resulting in p53 deletion and MYC overexpression in hepatocytes (frequent gene alterations in HCC patients) (**Fig. 1A**) (*32, 33*). Luciferase is expressed to enable *in vivo* tumor growth assessment (**Fig. 1B**), and we introduced a neoantigen in the form of membrane-anchored ovalbumin (mOVA) to monitor CD8, CD4 and B cell responses. The mOVA SIINFEKL epitope was mutated to the Y5 variant, to diminish H-2Kᵇ binding and CD8 T cell immunogenicity (*34, 35*), achieving 100% tumor growth (**Fig. 1C**). Early tumor lesions (d17) were classified as grade II, moderately differentiated HCC, and late-stage tumors (d28) classified as grade III, poorly differentiated HCC. In addition, late-stage tumors displayed high mitotic index and histopathological analysis revealed areas of necrosis and hemorrhage. These features indicate progressive tumor dedifferentiation and aggressive growth, establishing a model of advanced HCC in which immune regulation could be explored (**Fig. 1C,D**).

**Fig. 1.**
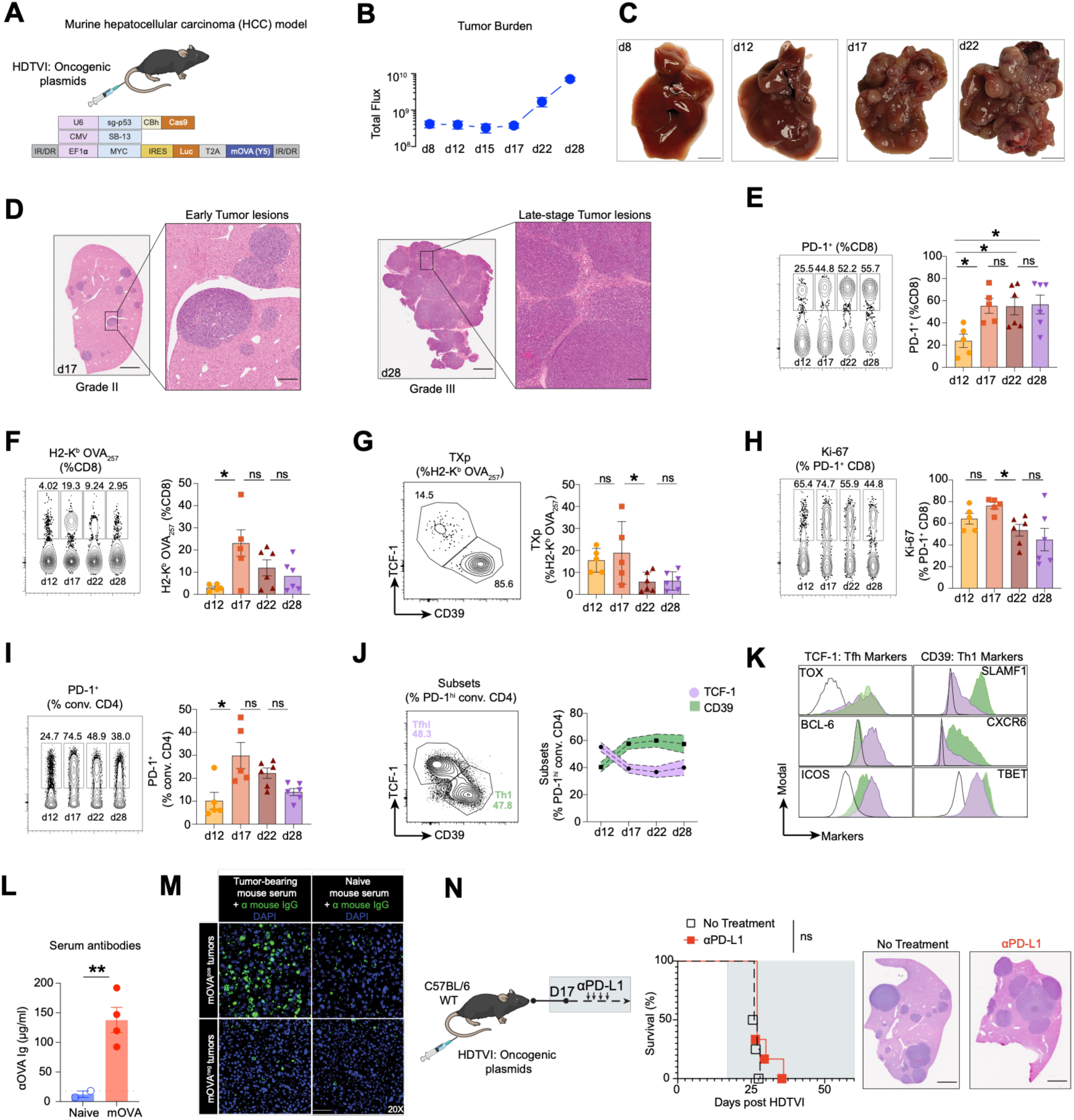
Autochthonous murine model of liver cancer resistant to immunotherapy despite CD8, CD4 and B cell responses. **(A)** Schematic of murine hepatocellular carcinoma (HCC) model: Hydrodynamic tail vein injection (HDTVI) was used to transfect hepatocytes and deliver oncogenic plasmids encoding: (1) single guide (sg) RNA targeting p53, as well as Cas9, (2) Sleeping Beauty transposase, (3) transposon based vector encoding MYC, luciferase (luc), and membrane ovalbumin (mOVA) containing SIIN<u>Y</u>EKL (Y5) mutation. **(B)** Longitudinal tumor burden measured by *In Vivo* Imaging System (IVIS) of luciferase activity and quantified by total flux. **(C)** Representative gross liver images at days (d) 8, 12, 17, and 22 after HDTVI. **(D)** Representative hematoxylin and eosin (H&E) stained liver sections of early and late-stage tumor lesions (17 and 28 days after HDTVI). Scale bars, 2.5 mm or 500 mm in insets. **(E)** Representative flow cytometry plots and graph show PD-1 expression among liver-infiltrating CD8 T cells at the indicated time points after HDTVI. **(F)** Representative flow cytometry plots and graph show H-2Kᵇ-OVA_257_ tetramer⁺ cells among liver-infiltrating CD8 T cells at the indicated time points after HDTVI. **(G)** Representative flow cytometry plot shows TCF-1 and CD39 expression among liver-infiltrating OVA-specific CD8 T cells. Graph shows frequency of TCF-1⁺CD39⁻ progenitor exhausted (TXp) at the indicated time points after HDTVI**.(H)** Representative flow cytometry plots and graph show Ki-67 expression among liver-infiltrating PD-1⁺ CD8 T cells at the indicated time points after HDTVI. **(I)** Representative flow cytometry plots and graph show PD-1 expression among liver-infiltrating conventional (Foxp3^-^) CD4 T cells at the indicated time points after HDTVI. **(J)** Representative flow cytometry plot shows TCF-1 and CD39 expression among liver-infiltrating PD-1^+^ conventional (conv) CD4 T cells. Graph shows TCF-1⁺CD39⁻ Tfhl and TCF-1⁻CD39⁺ Th1 subsets among PD-1⁺ conv. CD4 T cells at the indicated time points after HDTVI. **(K)** Representative histograms show expression of Tfh- and Th1-associated markers in TCF-1⁺ (purple) and CD39⁺(green) PD-1⁺ liver-infiltrating conventional CD4 T cells. Black lines show expression on naïve CD44^lo^ CD4 T cells. **(L)** Graph shows concentration of anti-OVA immunoglobulin (Ig) in serum of naïve mice and mice bearing mOVA tumors (22 days after HDTVI). (**M)** Representative immunofluorescence images show binding of serum IgG from mOVA tumor-bearing mice 22 days after HDTVI (left panels) or from naïve mice (right panels) to mOVA positive (top panels) or mOVA negative (bottom panels) HCC grown in RAGKO mice. Bound serum antibodies were detected with anti-mouse IgG, and counterstained with DAPI. Scale bars, 50 µm. **(N)** Experimental design shows αPD-L1 blockade treatment beginning on day 17 after HDTVI. Graph shows Kaplan–Meier survival curves for untreated and αPD-L1–treated mice. N= 4-6 per group. Right, representative H&E-stained liver sections from each group. Scale bars, 2.5 mm. Data in (E–I) are combined from two independent experiments and symbols represent individual mice. Data are shown as mean ± SEM. Data in (J and K) are representative from multiple experiments. Statistical significance was determined by one-way ANOVA with Tukey’s multiple comparisons test in (E, F, G, H, I) or Unpaired Welch’s t test in (L). For survival analysis (N), significance was determined by log-rank test. *P < 0.05, **P < 0.01, ns = non-significant.

Analysis of liver-infiltrating CD8 T cells revealed increased accumulation of PD-1^+^ CD8 T cells 17 days after HDTVI (**Fig. 1E and S1A**). Tumor-specific CD8 T cells identified by H-2K^b^-OVA257 tetramer staining peaked at day 17 (**Fig. 1F**). These were composed of more differentiated PD-1^+^ CD39^+^ exhausted cells (TX) as well as PD-1^+^ TCF-1^+^ progenitor exhausted CD8 T cells (TXp). TXp sustain PD-1^+^ CD8 T cell responses and are responsible for the proliferative burst of CD8 T cells that follows ICB (*36–39*). Indicative of progressive dysfunction, the frequency of tumor-specific TXp decreased with tumor growth (**Fig. 1G**) and we also observed a gradual reduction of proliferating PD-1^+^ CD8 T cells (**Fig. 1H**).

Similar to CD8 T cells, PD-1^+^ conventional CD4 T cells accumulated in the liver (**Fig. 1I**) and were comprised of TCF-1^+^ and CD39^+^ subsets (**Fig. 1J**). Although both populations expressed TOX, consistent with chronic stimulation, the TCF-1^+^ subset expressed markers associated with Tfh (*40–42*), whereas the CD39^+^ subset expressed markers associated with Th1 (*8, 43, 44*) (**Fig. 1K**). Regulatory CD4 T cell (Treg) frequencies increased by day 17 and remained similar through tumor progression (**Fig. S1B**). Assessment of additional immune compartments revealed increased infiltration of B cells at day 17 (**Fig. S1C**). Furthermore, we detected OVA-specific antibodies in the serum of tumor-bearing mice (**Fig. 1L**) and confirmed binding of these antibodies to mOVA-expressing HCC (**Fig. 1M**) and cells (**Fig. S1D**).

To determine whether PD-1/PD-L1 blockade was sufficient to control tumors in this model, mice with established HCC were treated with αPD-L1 blocking antibodies beginning on day 17 after tumor induction. Longitudinal assessment revealed no significant improvement in tumor control or survival following αPD-L1 treatment. Histological analysis of liver sections confirmed persistent tumor burden (**Fig. 1N**).

Together, these data support that this model resembles human tumors (*8*) with endogenous immune responses characterized by tumor-specific T cell activation and B cell responses, yet resistance to ICB.

### CD4 T cell-help sensitizes tumors to overcome resistance to PD-1 blockade

To determine whether enhancing tumor-specific CD4 T cells impacts tumor development, we adoptively transferred naïve OVA-specific TCR transgenic OTII CD4 T cells into mice 3 days after oncogenic plasmid delivery (Helped). Analysis at day 21 showed that OT-II CD4 T cells did not differentiate into Tregs and, consistent with chronic antigen stimulation, upregulated PD-1 and TOX when infiltrating the liver. Furthermore, similar to endogenous liver-infiltrating PD-1^hi^ CD4 T cells, OT-II differentiated into either TCF-1^+^ (Tfhl) or CD39^+^ (Th1) subsets (**Fig. 2A**). OT-II transfer was associated with dose-dependent reduction in tumor burden, demonstrating that tumor-specific CD4 T cells are beneficial for anti-tumor immunity in this model (**Fig. 2B**).

**Fig. 2.**
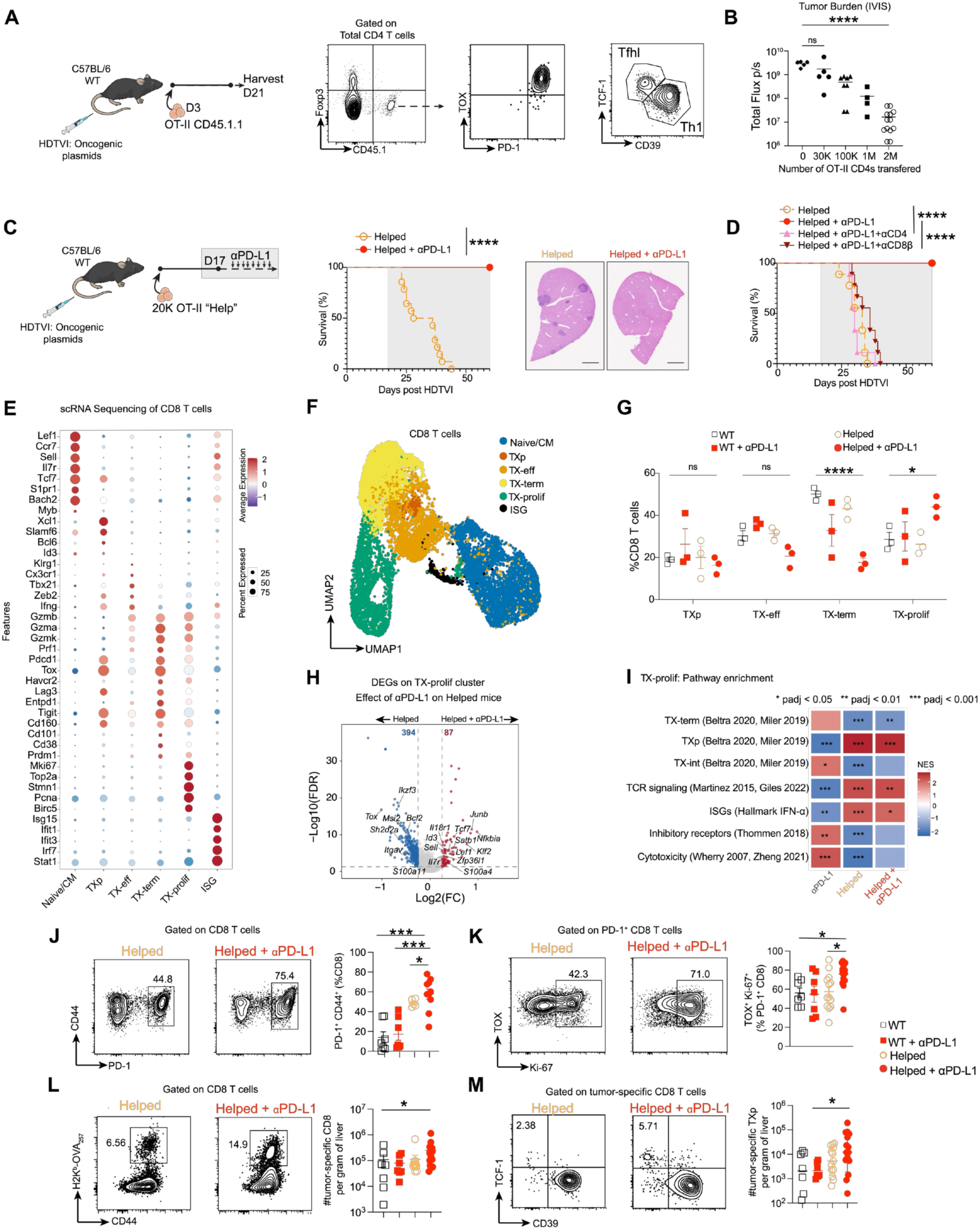
CD4 T cell-help sensitizes tumors to overcome resistance to PD-1 blockade. **(A)** Experimental design: Mice received hydrodynamic tail vein injection (HDTVI) of oncogenic plasmids on day 0, followed by adoptive transfer of CD45.1^+^ naïve OT-II CD4 T cells on day 3. Flow cytometry analysis of liver-infiltrating CD4 T cells was performed 21 days after HDTVI. Representative plots show expression of several markers on CD45.1^+^ OTII cells. **(B)** Tumor burden 21 days post HDTVI measured by IVIS of luciferase activity and quantified by total flux after transfer of the indicated numbers of OT-II CD4 T cells. **(C)** Experimental design: mice received 20,000 naïve OTII CD4 T cell transfer 3 days post HDTVI (Helped) and αPD-L1 treatment starting 17 days post HDTVI (Helped + αPD-L1). Graph shows Kaplan-Meier survival of Helped (N=14) vs Helped + αPD-L1 (N=12). Right, representative H&E-stained liver sections from each group. Scale bars, 2.5 mm**.(D)** Graph shows Kaplan-Meier survival curves. Mice received 20,000 naïve OTII CD4 T cell transfer (Helped) 3 days post HDTVI, αCD4 and αCD8β depletion antibodies on days 15 and 17, and αPD-L1 treatment starting day 17 post HDTVI, groups as indicated. N=9 per group. (E-M) Mice received 20,000 naïve OTII CD4 T cell transfer (Helped) 3 days post HDTVI and αPD-L1 treatment starting 17 days post HDTVI. Livers were collected for analysis 5 days post αPD-L1 treatment initiation (22 days post HDTVI). **(E)** Dot plots show gene expression (average and percent) of selected canonical markers across scRNA-seq–defined CD8 T cell clusters. **(F)** Uniform Manifold Approximation and Projection (UMAP) of intratumoral CD8 T cells (n = 9,707 cells) colored by annotated cluster identity: Naive/Central memory (CM), TXp (exhausted progenitor), TX-eff (exhausted effector-like), TX-term (exhausted terminal), TX-prolif (exhausted proliferating) and interferon-stimulated gene (ISG). **(G)** Quantification from scRNA-seq data of CD8 T cell subset proportions (percentage of total CD8 T cells) across treatment groups. N = 3 mice per group. **(H)** Volcano plot showing differentially expressed genes (DEGs) in the TX-proliferating cluster between Helped mice treated or not with αPD-L1. Numbers indicate DEGs at False Discovery Rate (FDR) < 0.05 and |log2FC| > 0.25. Red and blue denote upregulated and downregulated genes by αPD-L1 treatment, respectively. Select genes are labeled**.(I)** Gene set enrichment analysis (GSEA) of TX-prolif clusters against curated T cell differentiation and functional signatures. Heatmap displays normalized enrichment scores (NES) for each comparison. Gene sets derived from (*37, 58–61*) *Padj < 0.05, **Padj < 0.01, ***Padj < 0.001; Benjamini-Hochberg correction. **(J)** Representative flow cytometry plots and graph show frequency of PD-1^+^CD44^+^ cells among liver-infiltrating CD8 T cells. **(K)** Representative flow cytometry plots and graph show frequency of TOX^+^Ki67^+^ cells among liver-infiltrating PD-1^+^ CD8 T cells. **(L)** Representative flow cytometry plots show frequency and graph shows number per gram of liver of CD44^+^ H-2K^b^-OVA257 tetramer^+^ liver-infiltrating CD8 T cells. **(M)** Representative flow cytometry plots show TCF-1 and CD39 expression among tumor-specific (H-2K^b^-OVA tetramer^+^) CD8 T cells, and graph shows number of tumor-specific TXp (TCF-1^+^ cells) per gram of liver. Symbols represent individual mice. Data are shown as mean ± SEM. Statistical significance was determined by one-way ANOVA (B, J, K, L and M) or two-way ANOVA (G) with Tukey’s multiple comparisons test. For survival analysis in (C, D), significance was determined by log-rank test. ****P < 0.0001, ***P < 0.001, **P < 0.01, *P < 0.05; ns, not significant. Data in (J) are combined from 2 independent experiments and data in (K–M) are combined from 3 independent experiments.

Although CD4 T cell transfer modulated tumor burden, low-dose OT-II transfer was insufficient to confer durable tumor control. To model clinical observations and test whether augmenting CD4-help alters sensitivity to ICB, recipient mice that received low-dose OT-II (Helped) were treated with αPD-L1 (beginning at day 17 post HDTVI) (**Fig. 2C**). The combination of OT-II transfer and PD-L1 blockade resulted in complete tumor rejection and durable survival (**Fig. 2C, S2A**), demonstrating that the precursor frequency of tumor-specific CD4 T cells has a significant impact on the outcome to ICB.

To define the cellular requirements underlying responses to ICB response in mice receiving OT-II T cells, CD4 or CD8 T cells were depleted before αPD-L1 treatment. Depletion of either lineage abolished the survival benefit, resulting in tumor progression comparable to control group (**Fig. 2D**). Therefore, consistent with our data in patients (*8*), response to αPD-L1 requires the coordinated participation of both CD4 and CD8 T cells in this mouse model.

To assess how increased CD4 T cell-help and immunotherapy impacted the endogenous CD8 T cell compartment, we performed scRNA-seq on tumor-infiltrating immune cells (**Fig. S2B**). Subsetting of CD8 T cells revealed six transcriptionally distinct populations defined by canonical marker expression (**Fig. 2E, F and Fig. S2C**). Analysis of subset proportions revealed that the combination of CD4-help and αPD-L1 significantly expanded TX-prolif while reducing TX-term (**Fig. 2G**).

To further characterize transcriptional programs within the proliferating compartment, we performed differential gene expression analysis on TX-prolif cells across groups to evaluate the effect of αPD-L1 treatment and CD4-help (**Fig. 2H and S2E**). αPD-L1 induced only modest transcriptional changes in WT mice, whereas CD4-help substantially altered the TX-prolif response to αPD-L1. In Helped+αPD-L1 mice, TX-prolif cells showed increased expression of genes associated with progenitor cells, including *Il7r*, *Sell* (encoding CD62L), and *Tcf7* (encoding TCF-1) when compared to Helped mice (Fig. 2H). And TX-prolif cells in Helped+αPD-L1 mice showed increased expression of genes associated with activation and effector function (*Gzmb, Gzma, Ccl5, Ccl4, Jun, and Pim1*) when compared to untreated WT mice (**Fig. S2E**). These changes were accompanied by reduced expression of genes associated with inhibitory signaling, altered trafficking, and impaired function and survival including reduced *Tox*, *Sh2d2a*, *Bcl2, Ikzf3*. Consistent with this, pathway enrichment analysis on TX-prolif showed that the combination of CD4 T cell-help and αPD-L1 treatment was associated with enhanced progenitor-like, TCR signaling and interferon-associated programs, along with reduced terminal differentiation (**Fig. 2I**). Together, these data suggest that CD4-help qualitatively reshapes CD8 T cells responding to αPD-L1 therapy, expanding cells that retain progenitor-like features while acquiring effector activity and limiting terminal differentiation.

Flow cytometric analysis of liver-infiltrating cells at day 22 post HDTVI, confirmed that helped mice treated with αPD-L1 had increased frequency and proliferation of PD-1^+^CD44^+^ CD8 T cells (**Fig. 2J, K**). Help+αPD-L1 also significantly increased the number of H-2K^b^-OVA ^+^ tumor-specific CD8 T cells (**Fig. 2L**) and TXp (**Fig. 2M**) in the liver.

Together, these data demonstrate that tumor-specific CD4 T cells enhance expansion and quality of tumor-specific CD8 T cells during ICB, thus improving therapeutic outcome.

### Th1 and Tfh programs are both critical for efficacy of immunotherapy

To assess whether the therapeutic benefit conferred by tumor-specific CD4 T cells occurs during αPD-L1 treatment, we specifically depleted transferred Thy1.1^+^ OT-II cells prior to treatment initiation. Depletion of OT-II before treatment completely abrogated response to αPD-L1 (**Fig. 3A**), demonstrating that CD4-help during CD8 T cell priming is not sufficient for ICB efficacy and sustained tumor-specific CD4 T cell responses play a critical role in the success of immunotherapy.

**Fig. 3.**
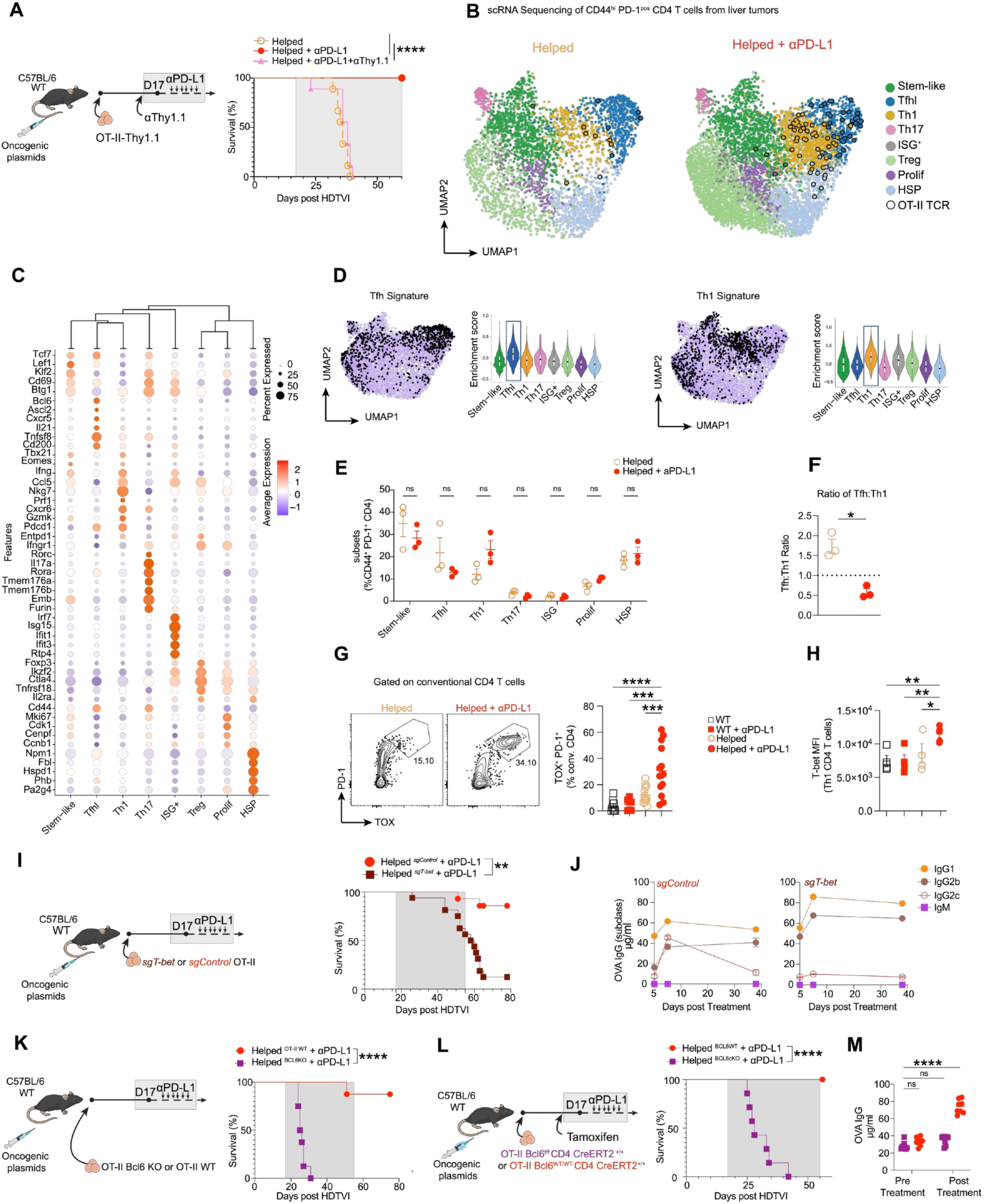
Th1 and Tfhl programs contribute to ICB efficacy. **(A)** Experimental design: mice received HDTVI on day 0 and 20,000 OT-II-Thy1.1 CD4 T cells on day 3, followed by αThy1.1 depleting antibody on days 15 and 17, and *a*PD-L1 therapy starting on day 17. Right: Kaplan-Meier survival curves. N=9 per group. **(B)** UMAP of liver-infiltrating PD-1^+^ CD4 T cells split by treatment condition and colored by cluster identity. Cells bearing the OT-II TCR (identified by paired TRAV14/TRBV12-1/TRBV12-2 CDR3 sequences) are outlined as open black circles.**(C)** Dot plots show gene expression (average and percent) of selected canonical markers across PD-1^+^ CD4 T cell clusters. **(D)** UMAP of Tfh (left) and Th1 (right) gene signatures and Violin plots of gene signature enrichment scores across PD-1^+^ CD4 T cell states.**(E)** CD44^+^PD-1^+^ conventional CD4 T cell subset proportions by scRNAseq. N=3 per group. Two-way ANOVA with Sidak’s multiple comparison test. ns = not significant; **(F)** Ratio of Tfh-like to Th1 CD4 T cells based on scRNAseq. Unpaired Welch’s t test *P < 0.05. **(G)** Representative flow cytometry plots and quantification of liver-infiltrating PD-1^hi^TOX⁺ conventional CD4⁺ T cells. One-way ANOVA with Tukey’s multiple comparisons test **P < 0.01. **(H)** T-bet mean fluorescence intensity (MFI) on PD1^hi^ CD39^+^ CD4 T cells (Th1). One-way ANOVA with Tukey’s multiple comparisons **P < 0.01. **(I)** Experimental design: Naïve OT-II CD4 T cells were electroporated with Cas9 and single guide (sg) RNA targeting F8 (*sgControl*) or *Tbx21* (*sgT-bet*) before adoptive transfer into recipient WT mice (20,000 OT-II CD4 T cells) that received HDTVI of oncogenic plasmids 3 days before. Mice received αPD-L1 treatment starting at day 17 post HDTVI. Right: Kaplan-Meier survival curves. Helped*^sgControl^* + αPD-L1 (N=14), Helped*^sgT-bet^* + αPD-L1 mice (N=16). Log-rank test. **P < 0.01. **(J)** OVA-specific serum IgG subclass (IgG1, IgG2b, IgG2c, and IgM) measured by ELISA over the course of αPD-L1 treatment. *sgControl* (n=6); *sgTbet (*n=8). Graph shows mean ± SEM.**(K)** Experimental design: OT-II *Bcl6*^f/f^ CD4-Cre-ER⁺ TdTomato⁺^/⁺^ mice received tamoxifen before isolation of TdTomato⁺ OT-II CD4 T cells (*Bcl6*KO) by cell sorting. A total of 20,000 *Bcl6*KO or WT OT-II CD4 T cells were transferred 3 days after HDTVI, and αPD-L1 treatment was initiated on day 17; Right: Kaplan-Meier survival curves. N=8 per group. Log-rank test. ****P < 0.0001. **(L)** Experimental design: Mice received HDTVI on day 0 and adoptive transfer of 20,000 *Bcl6*^fl/fl^ (*Bcl6*cKO) or *Bcl6*^wt/wt^ (*Bcl6*WT control) OT-II CD4-CreERT2^+/+^ TdTOM^+^/^+^ on day 3. Recipient mice received tamoxifen on day 15 and Day 16 followed by αPD-L1 treatment starting on day 17. Right: Kaplan-Meier survival curves. N=7 per group. Log-rank test. ****P < 0.0001. **(M)** OVA-specific serum IgG measured by ELISA 16 days post HDTVI (before αPD-L1) and 22 days post HDTVI (5 days after αPD-L1 treatment). Two-way ANOVA, ****P < 0.0001; ns = not significant. Each symbol represents one individual mouse, unless otherwise noted. Error bars represent mean ± SEM.

To define the effects of PD-L1 blockade on CD4 T cells, we performed paired single-cell RNA and TCR sequencing of liver-infiltrating CD44^+^ PD-1^+^ CD4 T cells 5 days after therapy initiation (**Fig. S3A**). Unsupervised clustering distinguished eight transcriptional states: Stem-like, Tfhl, Th1, Th17, interferon-stimulated gene (ISG)+, Treg, Proliferative and heat shock protein (HSP) (**Fig. 3B, C**). We identified transferred OT-II cells, by their canonical TCR α and β chain, and observed their segregation predominantly into Th1 and Tfhl states (**Fig. 3B**). scRNA sequencing also resolved a stem-like PD-1^+^ CD4 T cell subset, described in recent studies to sustain chronically stimulated CD4 T cells (*25, 45, 46*), which based on *Tcf7* (TCF-1) expression (**Fig. 3C**) was grouped with Tfhl on our flow cytometry analysis.

To validate the transcriptional identities of tumor-infiltrating PD-1^+^ CD4 T cells, we scored each cluster against published canonical gene signatures. Tfh signature scores were highest among the Tfhl cluster, while Th1 scores were higher in the Th1 cluster (**Fig. 3D**). Treg and cell cycle signatures also matched with their respective clusters (**Fig. S3B**). Cross-dataset comparison using published Tfh and Th1 signatures from viral infection with lymphocytic choriomeningitis virus (LCMV), vaccination, and multiple human cancer types confirmed that Tfhl and Th1 clusters aligned with bona fide Tfh and Th1 populations (**Fig. S3C**). Furthermore, these data confirmed that the stem-like PD-1^hi^ CD4 cluster was similar to self-renewing progenitor CD4 T cells that give rise to Tfh and effector Th1 present in chronic LCMV infection (*43, 45*) and in lymph nodes draining subcutaneous tumors (*45, 46*). Multidimensional scaling of CD4 subset transcriptional profiles further demonstrated that our murine HCC Tfhl cells clustered with canonical Tfh populations from LCMV and human tumors, while Th1 cells mapped closer to Th1/effector populations across different contexts (**Fig. S3D**).

Analysis of subset proportions within conventional CD44^hi^ PD-1^+^ endogenous CD4 T cells revealed no significant changes following *a*PD-L1 treatment (**Fig. 3E**), although the Tfhl:Th1 ratio was reduced following *a*PDL1 (**Fig. 3F**).

Flow cytometry analysis showed that *a*PD-L1 substantially increased the proportion of endogenous TOX^+^PD-1^+^ conventional CD4 T cells (**Fig. 3G**), but within this population there were no substantial changes in the relative frequency of CD39^+^ (Th1) subsets (**Fig**. **S3E**). However, consistent with the Tfhl:Th1 ratio shift observed by scRNA-sequencing, expression of Th1-defining transcriptional factor T-bet was increased in CD39^+^ PD-1^hi^ CD4 T cells in the liver of helped mice treated with *a*PD-L1, but not in TCF-1^+^ PD-1^hi^ CD4 T cells (**Fig. 3H and S3F**). Overall, these data support that both Tfh and Th1 tumor-specific CD4 T cells increase following PD-1 blockade, and consistent with previous reports, PD-1/PD-L1 blockade may favor Th1 differentiation (*47, 48*).

To better understand how PD-1 blockade modulates differentiation of PD-1^+^ CD4 T cells, we analyzed clonotypes that contained at least one stem-like cell and examined their distribution across stem-like, Tfh and Th1 states. In helped mice without ICB, these clones were confined to stem-like and Tfh states with little Th1 contribution, whereas in *a*PD-L1 treated helped mice there was a shift towards Th1 fate (**Fig. S3G**). Furthermore, we identified a clonotype present across all mice, and in helped mice, this public clone was restricted to stem-like and Tfh fates, whereas in helped + *a*PD-L1 mice, the same clone also gave rise to Th1 cells (**Fig. S3H**). In accordance with shared differentiation history between Tfhl and Th1 in tumors, we previously reported high clonal overlap between Tfhl (CXCL13^+^), Th1 and a CD4 T cell proliferating cluster (by Tversky asymmetric similarity index) in HCC patients that received neoadjuvant *a*PD-1 (*8*).

The segregation of tumor-specific CD4 T cells into Th1 and Tfh programs prompted us to test their relative importance for tumor control following ICB. To assess the contribution of Th1 differentiation, we evaluated the therapeutic benefit of OT-II CD4 T cells lacking the Th1 canonical transcription factor T-bet (generated by CRISPR-mediated deletion of *Tbx21*) (**Fig. 3I, S3I**). Mice that received T-bet-deficient tumor-specific CD4 T cells, presented much worse survival following *a*PD-L1 therapy than mice receiving WT tumor-specific CD4 T cells. Notably, whereas mice that received WT OT-II displayed a mixed anti-OVA IgG response (IgG1, IgG2c and IgG2b), mice that received T-bet-deficient OTII presented a marked reduction in IgG2c (**Fig. 3J**), consistent with the role of T-bet in promoting class-switch recombination to IFN-γ-associated isotypes (*49*).

To evaluate the contribution of Tfh differentiation, we used OT-II CD4 T cells isolated from *Bcl6*^fl/fl^ CD4-CreERT2 mice, in which tamoxifen administration results in deletion of the canonical Tfh transcription factor *Bcl6*. Donor mice were treated with tamoxifen before OTII isolation and transfer of *Bcl6*-deficient OTII completely abolished the help benefit to ICB response (**Fig. 3K**). In LCMV chronic infection *Xia et al,* reported that *Bcl6*-deficiency not only affected Tfh, but also the maintenance of Th1 effectors, due to *Bcl6*-dependency of stem-like CD4 progenitors (*45*). Therefore, given that *Bcl6* deficiency may prevent development of stem-like CD4 T cells in our HCC model, we performed a new experiment in which recipient mice received either *Bcl6*^fl/fl^ (*Bcl6* cKO) or *Bcl6*^WT/WT^ (*Bcl6* WT) CD4-CreERT2 OT-II cells and tamoxifen was only administered right before *a*PD-L1 therapy (**Fig. 3L**). Loss of *Bcl6* expression in OT-II cells, even after priming and differentiation, abolished the survival benefit conferred by ICB and abrogated the increase in OVA-specific IgG following αPD-L1 treatment (**Fig. 3M**). These findings demonstrate that sustained *Bcl6* expression in tumor-specific CD4 T cells is required for their helper function during ICB, while T-bet⁺ tumor-specific CD4 T cells also play a critical role in mediating ICB efficacy.

### Antigen-specific B cells are required for effective response to PD-1 blockade

Given the well-established requirement of BCL6 for Tfh and B cell help (*40, 41*), and the paramount role of BCL6 expression in helper CD4 T cells for production of tumor-specific IgG and therapeutic success of ICB, we sought to address the role of B cells and antibody-dependent cellular cytotoxicity (ADCC) mediated by NK cells. We depleted B cells (αCD20) or NK cells (αNK1.1) beginning two days before αPD-L1 treatment (**Fig. 4A**). Whereas NK cell depletion did not reduce the therapeutic efficacy of ICB, B cell depletion completely abolished the survival benefit. Flow cytometry analysis of tumor-infiltrating lymphocytes showed that depletion of B cells reduced BCL6 expression (**Fig. 4B**) as well as proliferation of TCF-1^+^ PD-1^hi^ conventional CD4 T cells (**Fig. 4C**), without affecting proliferation of the CD39^+^ Th1 population (**Fig. S4A**). These data indicate that B cells are required to sustain the Tfh program in tumors and is consistent with the well-known role of B cells in maintaining Tfh cells through antigen presentation and ICOS signaling in germinal centers (*23, 50*).

**Fig. 4.**
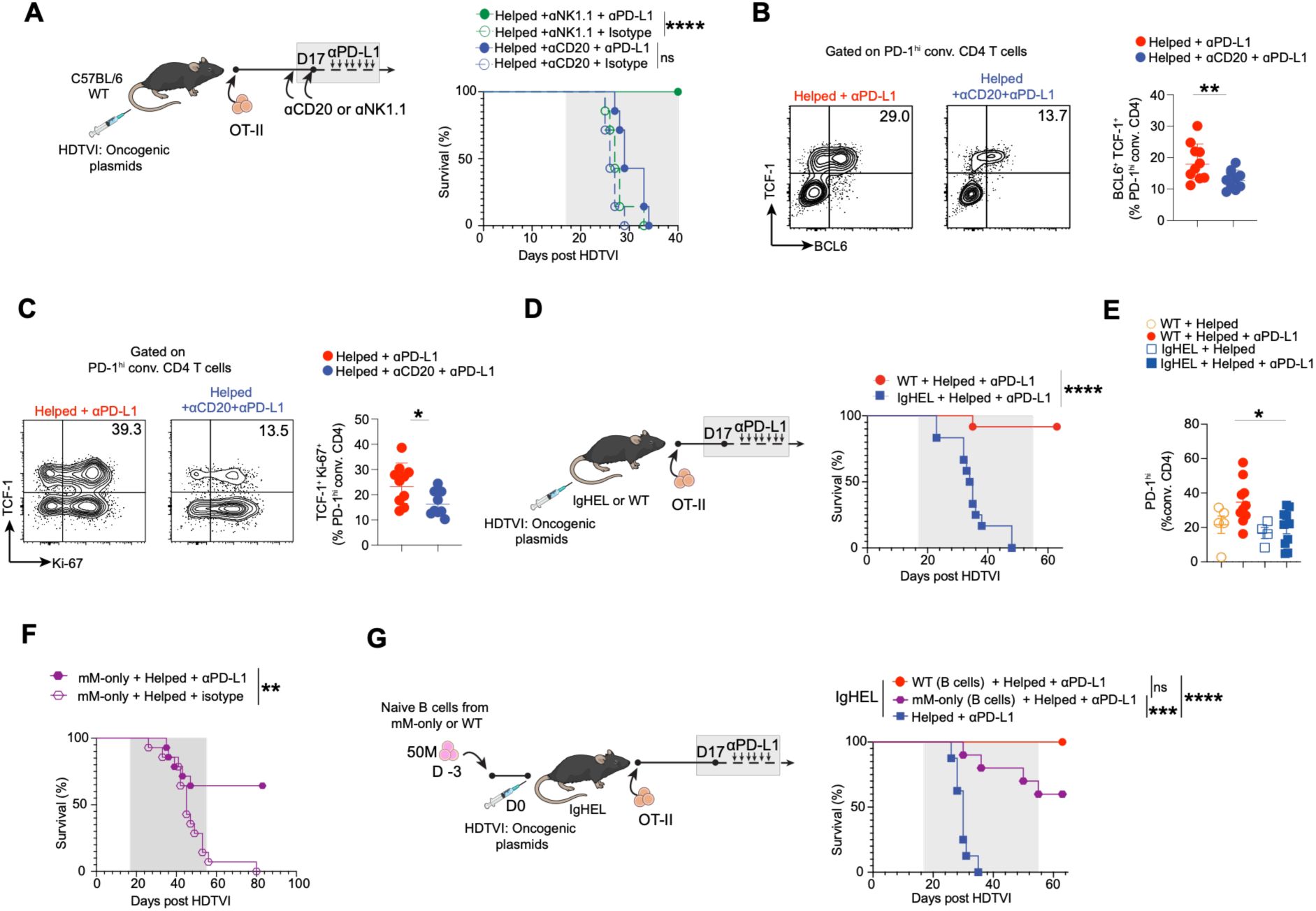
B cells - independently of secreted antibodies - support tumor-specific CD4 T cells for effective response to PD-1 blockade. **(A)** Experimental design: C57BL/6 mice received HDTVI on day 0 and 20,000 OT-II-CD4 T cells on day 3, followed by αCD20 (B cells) or αNK1.1 (NK cells) depleting antibodies on day 15 and 17, and αPD-L1 therapy or isotype control starting on day 17. Right: Kaplan–Meier survival curves. N=7 per group. Log-rank test. ****P < 0.0001. **(B)** Representative flow cytometry plots and graph show BCL6⁺TCF-1⁺ frequency among liver-infiltrating PD-1^hi^ conventional CD4⁺ T cells. Analysis was performed 5 days post αPD-L1 treatment initiation. Unpaired Welch’s t test. **P < 0.01. **(C)** Representative flow cytometry plots and graph show TCF-1⁺Ki-67⁺ frequency among liver-infiltrating PD-1^hi^ conventional CD4⁺ T cells. Analysis was performed 5 days post αPD-L1 treatment initiation. Unpaired Welch’s t test. *P < 0.05. **(D)** Experimental design: IgHEL transgenic mice or negative littermates (WT) received HDTVI on day 0 and 20,000 OT-II-CD4 T cells on day 3, followed by αPD-L1 therapy starting on day 17. Right: Kaplan–Meier survival curves. N=12 per group. Log-rank test. ****P < 0.0001 . **(E)** Graph shows frequency of liver-infiltrating PD-1^hi^ cells among conventional CD4⁺ T cells in WT or IgHEL mice receiving OT-II-help, treated with αPD-L1 therapy or not. Analysis was performed 22 days post HDTVI (5 days post αPD-L1 treatment initiation). One-way ANOVA with Tukey’s multiple comparison. *P < 0.05. **(F)** Kaplan–Meier survival curves of mM-only mice (*51*) that received 20,000 OT-II CD4⁺ T cells 3 days post HDTVI of oncogenic plasmids, and treated with αPD-L1 or isotype control starting 17 days post HDTVI. N=14 per group. Log-rank test. **P < 0.01. **(G)** Experimental design: IgHEL recipients received adoptive transfer of 50 x 10^6^ naïve splenic B cells from mM-only or WT donors 3 days before HDTVI of oncogenic plasmids. The control group did not receive B cell adoptive transfer. 3 days after HDTVI, all groups received 20,000 OTII CD4 T cells and starting 17 days post HDTVI, αPD-L1 treatment. Right: Kaplan–Meier survival curves. N= 8-10 per group. Pairwise survival comparisons were performed using log-rank tests. ****P < 0.0001, ***P < 0.001. ns = not significant. Each symbol represents one individual mouse, unless otherwise noted. Error bars represent mean ± SEM.

To determine whether the B cell contribution is antigen-specific, we used IgHEL transgenic mice, in which B cells express a singular BCR specific for hen egg lysozyme (HEL) that does not cross-react with OVA. Transfer of OT-II cells into IgHEL hosts followed by αPD-L1 treatment resulted in complete loss of survival benefit, compared to WT hosts (**Fig. 4D**). Flow cytometry analysis of liver-infiltrating cells revealed that OTII cells failed to expand following PD-L1 blockade in IgHEL mice (**Fig. S4B**). IgHEL hosts also showed impaired increase in endogenous liver-infiltrating PD-1^hi^ CD44^+^ conventional CD4 T cells after immunotherapy (**Fig. 4E**), indicating that antigen-specific B cells are required for sustaining CD4 T cell responses within the tumor. Overall, these data show that antigen-specificity of B cells is critical for therapeutic efficacy of ICB, consistent with an important role for cognate interactions between B cells and helper CD4 T cells.

To assess the role of secreted Ig in the response to ICB, we used mice engineered to express only membrane-bound IgM (mM-only) and no secreted antibodies (*51*). mM-only B cells express a normal BCR repertoire and retain the ability to capture and present antigens to sustain Tfh cells and develop germinal centers (*51*). mM-only mice that received OT-II transfer and αPD-L1 showed significant survival benefit compared to isotype-treated controls (**Fig. 4F**), demonstrating that antibody secretion is not required for therapeutic response to ICB. Finally, we directly compared the ability of mM-only and WT B cells in rescuing response in IgHEL hosts by adoptive transfer of naïve B cells before tumor induction and OT-II transfer (**Fig. 4G**). Transfer of WT B cells fully rescued the survival benefit of IgHEL hosts, whereas mM-only B cells provided substantial rescue compared to IgHEL hosts without B cell reconstitution (**Fig. 4G**). These data demonstrate that the prominent role of B cells in ICB is not dependent on antibody secretion. These data parallel germinal center biology, where maintenance of the Tfh cell phenotype requires sustained antigenic stimulation by germinal center B cells, suggesting that B cells may similarly support tumor-specific CD4 T cells through antigen-dependent interactions (*23, 52*).

## DISCUSSION

Our previous analysis of HCC patients that received PD-1 blockade highlighted the role of Tfhl cells and B cells in tumor control (*8, 28*). CD4 T cell-help is increasingly recognized as a determinant of ICB response (*7–9, 25, 53, 54*), yet the specific contributions of CD4 T cells and the role of B cells, remain undefined. In this study, we demonstrate that coordinated responses of CD8, CD4 and B cells are essential for long-term HCC survival following PD-L1 blockade. We show that tumor-specific Th1 and Tfh cells are critical and uncover the essential antibody-independent cognate-dependent function of B cells following ICB.

Similar to increased intratumoral frequency of CD4-helpers in HCC patients that respond to PD-1 blockade, here we show that increasing the precursor frequency of tumor-specific CD4 T cells sensitized HCC tumors to PD-L1 blockade. Help + ICB augmented tumor-specific CD8 T cells, including TXp, the subset that maintains PD-1^+^ CD8 T cell responses and provides the proliferation burst in response to PD-1/PD-L1 blockade (*36–38*). Responding CD8 T cells proliferating after ICB exhibited features of a less terminally differentiated state, consistent with the maintenance of durable effector potential and long-term tumor control. PD-L1 blockade also drove the expansion of tumor-specific CD4 T cells, augmenting both Tfh-like and Th1 compartments while selectively skewing dominant clones toward a Th1 phenotype, as previously described (*47, 48*). Nevertheless, antitumor immunity depended on the coordinated activity of both CD4 T-cell programs. Genetic disruption of either *Bcl6* or T-bet on transferred tumor-specific CD4 helpers abrogated therapeutic benefit and eliminated long-term survival, establishing Tfh-like and Th1 states as complementary and indispensable components of effective checkpoint responses. Importantly, inducible deletion of *Bcl6* on tumor-specific CD4 helpers at treatment initiation also abolished tumor control, revealing a continuous requirement for Tfh-like CD4 T cells beyond initial priming. Consistent with recent studies, clonal analysis linked the Tfhl and Th1 states through a shared stem-like CD4 population, paralleling the *Bcl6*-dependent progenitor described in chronic viral infection (*45*), with corresponding Tfhl–Th1 clonal overlap in human HCC (*8*). A plausible division of labor assigns the *Bcl6*-dependent Tfhl arm to maintenance of chronically stimulated CD4 T cells, and the T-bet-dependent Th1 arm to inflammatory functions that condition APCs and support CD8 T cell immunity.

In this model, we detected antibodies to membrane OVA expressed on transformed hepatocytes, and anti-OVA IgG increased following PD-L1 blockade. The surge in anti-OVA IgG was dependent on *Bcl6* expression on helper OT-II cells, and IgG2c (similar effector activity to human IgG1)(*55*) required T-bet-expressing OT-II cells. Yet mM-only mice, which are incapable of secreting antibodies, retained responsiveness to ICB and exhibited improved survival compared to isotype control treated animals. These findings reveal a dominant secreted antibody-independent role for B cells in orchestrating effective antitumor immunity. These observations do not preclude that secreted antibodies may play a role in other tumors or when directed to other antigens (*56*). However, these data suggest that the association between IgG1 signatures and clinical response to checkpoint blockade may reflect ongoing antigen-specific CD4–B cell interactions rather than a direct effector function of secreted antibodies. Consistent with a critical role for antigen-specific B-cell responses, IgHEL transgenic hosts failed to support the expansion of PD-1^hi^ conventional CD4 T cells following PD-L1 blockade, and without B cell cognate interactions, IgHEL mice derived no survival benefit from ICB. Likewise, B-cell depletion markedly reduced the frequency of PD-1^hi^ TCF-1⁺BCL6⁺ Tfh-like cells and abrogated the therapeutic efficacy of ICB. Based on these data we propose that B cells play a major sustaining tumor-specific CD4 helper T cells. Similar to interactions in germinal center reactions (*23, 57*), we propose that antigen presentation by B cells enforces the Tfh program to endow CD4 T cells to survive with persistent antigen stimulation.

Collectively, our data support a model in which durable responses to checkpoint blockade require coordinated adaptive immune networks involving CD8 T cells, CD4 T cells, and antigen-specific B cells. Rather than acting as independent effectors, these populations form an interdependent immune circuit that sustains productive antitumor immunity and enables long-term tumor control.

## Acknowledgments

We thank the Center for Comparative Medicine and Surgery, Dean’s Flow Cytometry CORE, Biomedical Engineering and Imaging Institute, and other Core facilities at the Icahn School of Medicine at Mount Sinai. We thank Zhihong Chen and Rachel Chen from the Human Immune Monitoring Center at the Icahn School of Medicine at Mount Sinai for their help with single-cell RNA sequencing. We thank the NIH Tetramer Core Facility (NIH Contract 75N93020D00005 and RRID:SCR_026557) for providing H2K^b^-OVA257 monomers. We thank Kai-Hui Yao for her help with experiments with mM-only mice. Computational work was performed on Minerva HPC and was supported in part through the computational and data resources and staff expertise provided by Scientific Computing and Data at the Icahn School of Medicine at Mount Sinai.

## Author contributions

Conceptualization: AOK, AV Methodology: AOK, AV, AL

Investigation: AV, EH, NP, V.v.H, LB, JH, RM, AL, PH, BC, SG, IK, KEL, SS, HH, GF and AOK

Computational: AV, NP, EGK, BRR, AOK Visualization: AV, AOK, V.v.H

Funding acquisition: AOK, MCN Project administration: AOK

Supervision: AOK, MCN, MM, AL, SG, BRR

Writing – original draft: AV, AOK

Writing – review & editing: All authors.

## Data, code, and materials availability

No New software pipelines were used in the study beyond those described in relevant Methods sections. Single-cell RNA-seq and cell-hashing data for CD45 sorted dataset have been deposited in GEO# GSE337198 and Single-cell RNA-seq, TCR-seq, cell-hashing data have been deposited in GEO#GSE337195 and will be made available at the time of publication.

## Dissertation statement

The data in this paper were used in a dissertation as partial fulfillment of the requirements for a PhD degree at the Graduate School of Biomedical Sciences at the Icahn School of Medicine at Mount Sinai, A.V.

