## Supplementary Files for "B cells sustain tumor-specific CD4 T cells to promote response to PD-1 targeted therapy"

**The PDF file includes:**

Materials and Methods

Figs. S1 to S4

#### MATERIALS AND METHODS:

##### Mice

Mice were used as donors for immune cells or injected with oncogenic plasmids at 7–8 weeks of age or at least 18 g of weight; males and females were used. C57BL/6JOlaHSd (WT) mice were purchased from Inotiv (previously Envigo Labs). For naïve CD4 transfers, OT-II TCR transgenic mice (strain 004194) were purchased from Jackson Laboratory and bred with CD45.1 (strain 002014, B6.SJL-Ptprca Pepcb/BoyJ) or Thy1.1 (strain 000406). IgHEL-MD4 (C57BL/6-Tg(IghelMD4)4Ccg/J) mice were purchased from Jackson Laboratory (strain 002595) and bred in house to C57BL/6JOlaHSd to obtain IgHEL transgenic and WT littermates for experiments. To match the microbiome with WT mice from Inotiv, bedding from incoming Inotiv mice was transferred into IgHEL breeding cages regularly. mM-only mice were generated and maintained at The Rockefeller University by the Nussenzweig lab (*51*). For *Bcl6* disruption in OT-II CD4 T cells, *Bcl6*$^{fl/fl}$ CD4-CreERT2 TdTomato $^{+}$/$^{+}$ mice (a kind gift from Dr. Maria Lafaille, ISMMS) were bred with OT-II CD45.1.1 mice. RAG-KO mice were a kind gift from Dr. Glaucia Furtado and were maintained at the Icahn School of Medicine at Mount Sinai. All animal experiments were approved by the Institutional Animal Care and Use Committee of the Icahn School of Medicine at Mount Sinai (protocol IACUC-2018-0018 / PROTO201900610) and by the Rockefeller University Institutional Animal Care and Use Committee (IACUC-24020-H).

##### Naïve CD4 T cell electroporation and adoptive T cell transfer

sgRNAs targeting murine T-bet (sgT-bet_1: AGUCUGGGUGGACAUAUAAG; sgT-bet_2: AGGACUACGCAUUGCCCGCG) and F8 (used as a control, not expressed in T cells; sgF8: TCTCCAATGAAGCTTGACGG) were purchased from Synthego (Gene Knockout Kit, previously CRISPR Revolution sgRNA EZ Kit) and guides were selected from Brie

CRISPR library (*73*). For sgRNA/Cas9 RNP formation, 1 µl of sgRNA (0.3 nmol/µl in nuclease-free H2O) was incubated with 0.6 µl of Cas9 (10 mg/ml, Alt-R S.p. Cas9 Nuclease V3, Integrated DNA Technologies) in nuclease-free H2O in a final volume of 5 µl for 10 min at room temperature. A combination of both guides was used (0.5 µl of each gRNA incubated with Cas9 for 10 min).

CD4 T cells were enriched from splenocytes of naïve OT-II mice with the MojoSort Mouse CD4 T Cell Isolation Kit (BioLegend, 480033) according to the manufacturer's instructions. $2 \times 10^6$ enriched OT-II CD4 T cells were washed in PBS, resuspended in 20 µl P3 buffer (P3 Primary Cell 4D-Nucleofector X Kit S, Lonza), mixed with the 5 µl sgRNA/Cas9 RNP, and immediately electroporated using the 4D-Nucleofector System (pulse DS-137)(*15*). Immediately after electroporation, 150 µl of media (RPMI, 10% FBS, 2 mM L-glutamine, 10 mM HEPES, 55 µM β-mercaptoethanol) was added to each well, and cells were rested for 1 h at 37 °C before further processing (Nüssing et al., 2020; Seki and Rutz, 2018). 20,000 OT-II CD4 T cells were transferred i.v. into tumor-recipient mice 3 days post-hydrodynamic injection of oncogenic plasmids.

For transfer of *Bcl6*-disrupted OT-II cells, *Bcl6*$^{fl/fl}$ CD4-CreERT2 TdTomato $^{+}/^{+}$ donor mice received tamoxifen (Sigma, T5648) dissolved in corn oil (Sigma, C8267) at 5 mg per dose (100 µL) by oral gavage on two consecutive days. Two days after the final dose, spleens were harvested, CD4 T cells were enriched by magnetic selection (BioLegend, 480033), and TdTomato$^{+}$ cells were FACS-sorted. $2 \times 10^4$ sorted CD4 T cells were transferred i.v. into recipient mice 3 days post-HDTVI.

For B cell isolation, spleens were collected from C57BL/6 (WT) or mM-only mice, disrupted through 100 µm strainers, treated with RBC lysis buffer (ACK, Gibco) for 3 min at room temperature, and enriched using the MojoSort Mouse Pan B Cell Isolation Kit (BioLegend, 480051) according to the manufacturer's instructions. $5 \times 10^7$ naïve B cells were transferred i.v. into IgHEL recipient mice 3 days prior to HDTVI of oncogenic plasmids.

**Hydrodynamic injection of oncogenic plasmids**

A sterile saline/plasmid mix was prepared containing 10 µg of pT3-EF1a-MYC-IRES-mOVA (mOVA), 20 µg of px330-sg-p53 (sgp53), and 2.5 µg of SB13 transposase-encoding plasmid (kind gift from Dr. Xin Chen, University of Hawaii Cancer Center) dissolved in 2 ml of saline; 10% of each mouse's body weight was injected in volume(*74*). Vectors for hydrodynamic delivery were produced in house (Qiagen, 12362) or by Azenta Life Sciences. Equivalent DNA concentration between batches was confirmed by gel equivalence to ensure reproducibility.

#### Vector design

pT3-EF1a-MYC-IRES-mOVA was generated by cloning mOVA from pCI-neo-mOVA (gift from Maria Castro, Addgene #25099) into pT3-EF1a-MYC-IRES-lucOS (Ruiz de Galarreta et al., 2019; Addgene #129776). The SIINFEKL epitope ($OVA_{257}$) was mutated to SIINYEKL (Y5) to reduce MHC-I binding affinity (*34, 35*). Vector maps and plasmids will be deposited on Addgene upon publication. pT3-EF1a-MYC-IRES-Luc plasmids were generated by the lujambio lab (Addgene#129775)

#### TIL isolation from liver tumors

Liver was weighed after collection and approximately 1 g was used for processing. Liver was minced and digested in 15 ml of collagenase IV (Sigma, C5138; 0.25 mg/ml) and DNase I (Sigma, DN25; 0.1 mg/ml) for 30 min at 37 °C. Tissue was further homogenized with a 10 ml syringe and 16G needle, filtered through a 100 µm strainer, washed with media, and filtered again through a 70 µm strainer. Cells were resuspended in 25% Percoll (Cytiva, 17089101), underlaid with 70% Percoll, and spun at 400 g for 20 min without acceleration or brake. The lymphocyte interface was collected with a transfer pipette. RBC lysis was performed with ACK buffer (Gibco) for 3 min at room temperature. Cells were stained immediately unless assessed for intracellular proteins; in vitro stimulation was performed with PMA/ionomycin (BioLegend, 423301) for 3 h at 37 °C. Phenotypic analysis of TILs were performed at day 22 post HDTVI.

#### In vivo treatments and antibody-mediated depletions

For CD4 and CD8β depletion, mice received 200 µg each of anti-CD4 (Bio X Cell, clone GK1.5, BE0003-1) and anti-CD8β (Bio X Cell, clone 53-5.8, BE0223) i.p. on days 15 and 17 post-HDTVI. For Thy1.1 depletion, mice received 200 µg anti-Thy1.1 (Bio X Cell, clone 19E12, BE0214) on days 15 and 17 post-HDTVI. For B cell and NK cell depletion, mice received 200 µg anti-CD20 (Bio X Cell, clone MB20-11, mouse IgG2c,κ, BE0356) or 200 µg anti-NK1.1 (Bio X Cell, clone PK136, BE0036) on days 15 and 17 post-HDTVI. For checkpoint blockade, mice received 200 µg rat IgG2b isotype control (Bio X Cell, clone LTF-2, BE0090) or 200 µg anti-PD-L1 (Bio X Cell, clone 10F.9G2, BE0101) i.p. every 3 days starting on day 17 post-HDTVI.

#### Antibodies and flow cytometry

Single-cell suspensions were surface-stained for 30 min on ice in flow cytometry buffer (PBS, 2% FBS, 1 mM EDTA, 0.05% sodium azide) supplemented with 10% Brilliant Stain Buffer (BD Biosciences) if more than two Brilliant Violet dyes were used, and with anti-CD16/32 (TruStain FcX, BioLegend). OVA-derived H-2K$^b$/OVA$_{257}$ biotinylated monomers were obtained from the NIH Tetramer Core Facility and tetramerized as previously described (Altman et al., 1996); tetramer staining was performed before surface antibody staining. Cells were incubated with fixable viability dye (Thermo Fisher) in PBS for 10 min at room temperature. For intracellular cytokine staining, samples were fixed/permeabilized overnight at 4 °C with the Foxp3/Transcription Factor Staining Buffer Set (eBioscience), and intracellular staining was performed at room temperature for 45 min.

Antibodies used for spectral flow cytometry were as follows: Ki67 (BUV395, clone B56, BD, 564071), CD4 (BUV496, clone GK1.5, BD, 612952), CD8a (BUV737, clone 53-6.7, BD, 612759), CD39 (BUV805, clone 24DMS1, BD, 368-0391-80), TCF-1 (Pacific Blue, clone C63D9, Cell Signaling Technology, 9066S), CD45.2 (BV510, clone 104, BioLegend, 109838), B220 (BV605, clone RA3-6B2, BioLegend, 103243), CD19 (BV650, clone 1D3, BD, 563235), CXCR6 (BV711, clone SA051D1, BioLegend, 151111), SLAMF1/CD150 (BV711, clone TC15-12F12.2, BioLegend, 115941), PD-1 (BV785, clone 29F.1A12, BioLegend, 135225), TOX (Vio515, clone REA473, Miltenyi Biotec, 130-129-208), *Bcl6* (PE, clone IG191E/A8, BioLegend, 648304), ICOS (PE-Cy5, clone 15F9, BioLegend, 107708), T-bet (PE-Cy7, clone 4B10, BioLegend, 644823), Foxp3 (eFluor 660, clone FJK-16s, Invitrogen, 50-5773-82), CD44 (AF700, clone IM7, BioLegend, 103026), CD45.1 (APC-Cy7, clone A20, BioLegend, 110716), and CD3/CD3e (APC-Fire 810, clone 145-2C11, BioLegend, 100312). Antigen-specific CD8 T cells were identified using H-2K$^b$/OVA$_{257}$ (SIINFEKL) tetramer, generated in house by tetramerization of biotinylated monomers obtained from the NIH Tetramer Core Facility.

#### ELISA

Semi-quantitative ELISA for antigen-specific mouse antibodies was performed as follows. High-binding 96-well plates (Nunc MaxiSorp) were coated overnight at 4 °C with OVA (2 µg/mL) in PBS; standard wells were direct-coated with a dilution series of purified mouse immunoglobulin matching the isotype being measured. Plates were washed in PBST (PBS + 0.05% Tween-20) and blocked with 1–3% BSA in PBS for 1 h at room temperature. Serum was applied at 1:25, 1:50, and 1:100 for 1 h at room temperature, and bound antibody was detected with HRP-conjugated anti-mouse IgG1, IgG2b, IgG2c, or IgM (and other isotype-specific HRP secondaries as needed; 1:5000, 30–60 min).

Wells were developed with TMB, stopped with 1 N $H_2SO_4$, and read at 450 nm (reference 570–620 nm).

#### Bioluminescence (luciferase) imaging

In vivo bioluminescence imaging was performed using an IVIS Spectrum system (Caliper Life Sciences) to quantify liver tumor burden before mice were evenly assigned to treatment cohorts. Mice were imaged 5 min after intraperitoneal injection of fresh D-luciferin (150 mg/kg; Thermo Scientific). Luciferase signal was quantified using Living Image software (Caliper Life Sciences). Normalized luciferase flux was calculated by subtracting background signal. Each treatment cohort had equivalent average luciferase flux.

#### Immunofluorescence (IF) detection of serum antibodies bound to tumor tissue

$mOVA^+$ liver tumors were induced by hydrodynamic tail vein injection of a MYC-Luc-mOVA plasmid or control mOVA− tumors with a MYC-Luc plasmid (*33*), into Rag1−/− mice; Rag1-/- were a kind gift from Dr. Glaucia Furtado. Rag1−/− hosts were used to eliminate endogenous immunoglobulin background. Tumor-bearing liver sections were incubated with serum from tumor-bearing or naive mice as the primary antibody source, and bound mouse antibody was detected with a goat anti-mouse IgG secondary antibody conjugated to Alexa Fluor 647 (H+L, highly cross-adsorbed; Invitrogen, A-21236). Nuclei were counterstained with DAPI. Whole-slide fluorescence images were acquired at 20× with identical settings across all conditions.

#### Flow cytometric serum antibody binding assay.

HEK-293T cells were transfected with the MYC-Luc-mOVA (mOVApos) or control MYC-Luc (mOVAneg) plasmid using Lipofectamine 3000 (Invitrogen). Three days post-transfection, HEK cells were harvested and stained with serum from mOVA tumor-bearing mice (1:10, 1 h, on ice), and bound mouse IgG was detected with donkey anti-mouse IgG (H+L) Alexa Fluor 594.

#### Single-cell RNA sequencing

C57BL/6 mice were injected with oncogenic plasmids by HDTVI. Three days post-HDTVI, 20,000 naïve OT-II CD4 T cells were adoptively transferred i.v. Treatment groups were stratified by tumor burden (IVIS), treatment was initiated on day 17, and mice were euthanized on day 22. Livers were harvested and processed to single-cell suspensions, stained for CD45 (PE) and Live/Dead IR for 30 min on ice, washed three times (PBS, 1 mM EDTA, 2% FBS), and $CD45^+$ TILs were sorted as live $CD45^+$ by FACS (BD FACSAria). Viability of FACS-sorted cells was assessed by Acridine Orange/Propidium Iodide staining (Nexcelom), showing 97% viability. Sorted $CD45^+$ cells were then labeled with a panel of 12 unique TotalSeq-C hashtag oligonucleotide (HTO)-conjugated antibodies (BioLegend, #155801–155812) to enable sample multiplexing and identification of individual mice. Cells from all mice were pooled in equal proportions for scRNA-seq processing.

#### $CD45^+$ intratumoral immune cell library preparation and sequencing

scRNA-seq was performed on the Chromium GEM-X platform (10x Genomics) using the GEM-X Single Cell 5′ gene expression (5′ GEX) v3 kit, targeting recovery of 50,000 cells per lane. GEMs were generated on the sample chip in the Chromium X system. Barcoded cDNA was extracted after post-GEM RT cleanup and amplified for 12 cycles. Full-length cDNA underwent enzymatic fragmentation, end-repair, A-tailing, adapter ligation, and sample-index PCR for library construction per the 10x Genomics Chromium Single Cell 5′ Reagent Kits v3 User Guide (CG000331). Twelve samples (4 conditions × 3 biological replicates) were multiplexed using HTOs. Libraries were quantified by TapeStation (Agilent) and Qubit (ThermoFisher) and sequenced paired-end on the NovaSeq X (Illumina), targeting 20,000 reads per cell for gene expression and 5,000 reads per cell for HTO across two lanes. Raw fastq files were aligned to the mm10 reference genome (2020-A) and demultiplexed using Cell Ranger multi v7.0.1.

#### $CD44^+PD\text{-}1^+$ CD4 T cell scRNA-seq and TCR-seq library preparation and sequencing

$CD44^+PD\text{-}1^+$ CD4 T cells were FACS-sorted and processed for concurrent 5′ gene expression, HTO (feature barcode), and V(D)J TCR libraries using the Chromium GEM-X Single Cell 5′ v3 kit and the Single Cell V(D)J Enrichment Kit for mouse T Cells (10x Genomics). Six samples (3 replicates × 2 conditions: OT-II and OT-II + αPD-L1) were multiplexed with 6 TotalSeq-C hashtag antibodies (BioLegend; HTO-2/3/9 = OT-II; HTO-4/5/6 = OT-II + αPD-L1) and loaded as a single pool. GEMs were generated on the sample chip in the Chromium X system. Barcoded cDNA was amplified for 12 cycles and split for gene expression library construction, TCR V(D)J targeted enrichment with locus-specific primers (Chromium Single Cell V(D)J Reagent Kits User Guide, CG000206), and Feature Barcode (HTO) library construction (CG000208). Libraries were quantified by TapeStation (Agilent) and Qubit (ThermoFisher) and sequenced paired-end on the NovaSeq X, targeting 20,000 reads per cell for gene expression and 5,000 reads per cell for TCR and

HTO. Raw fastq files were aligned to mm10 (2020-A) and demultiplexed using Cell Ranger multi v7.0.1, which simultaneously processed GEX, HTO, and VDJ libraries.

#### Single-cell RNA-seq analysis

##### CD45$^+$ cells

Downstream analysis was performed in R (v4.2.0) using Seurat v5 (Hao et al., 2024). Raw count matrices from two sequencing pools (12 samples each; 4 conditions × 3 replicates) were loaded from Cell Ranger multi output. Cells were filtered (200–6,000 genes, <50,000 UMIs, <10% mitochondrial content), and doublets were removed using scDblFinder (Germain et al., 2021), retaining 55,768 cells. Each pool was normalized with SCTransform (v2) and pools were integrated using SCT-based anchor integration with 3,000 variable features. PCA (50 PCs), UMAP, and Louvain clustering (dims 1:30, resolution 0.5) were performed on the integrated assay. Clusters were annotated into broad immune populations using canonical markers, then each compartment was subset and re-clustered with resolution chosen via Clustree (v0.5.1; Zappia and Oshlack, 2018). For the CD8 subset (9,707 cells), cell-cycle scores were regressed out using SCTransform (vars.to.regress = c("S.Score", "G2M.Score")) prior to re-clustering (resolution 0.5), yielding Naïve/CM, TXp, TX-eff, TX-term, TX-prolif, and ISG populations. Cluster marker genes were identified with FindMarkers (Wilcoxon). Between-condition differential expression was performed on the RNA assay using MAST (v1.30.0) fit as a two-part hurdle model with a random effect for sample (MAST-RE; zlm, method = "glmer"), with the cellular detection rate included as a covariate, to account for within-mouse correlation among cells (Zimmerman et al., 2021; FDR < 0.05, |log2FC| > 0.25). Condition-level differences were additionally assessed by pseudobulk aggregation of sample-level raw counts with limma-voom, as an orthogonal replicate-level test. Single-cell RNA-seq and cell-hashing data for the CD45-sorted dataset have been deposited in GEO# GSE337198.

##### Sorted CD44$^+$PD-1$^+$ CD4 T cells

Six HTO-multiplexed samples (3 replicates × 2 conditions; HTO-2/3/9 = OT-II, HTO-4/5/6 = OT-II + αPD-L1) were loaded from Cell Ranger multi output. Cells were filtered (200–6,000 genes, <15% mitochondrial content), demultiplexed with HTODemux (positive.quantile = 0.99), and singlets retained (9,772 cells). SCTransform (v2, regressing percent.mt) was followed by PCA, UMAP, and clustering (dims 1:30). Resolution was selected via Clustree (v0.5.1), and a resolution of 0.8 was used for final clustering, yielding nine clusters (Stem-like, Tfhl, Th1, Treg, Th17, Proliferating, ISG, HSP and Effector). Clusters were annotated by canonical markers and module scoring (Seurat

AddModuleScore, default parameters; Tfh: Cxcr5, Icos, Bcl6, Tcf7, Pdcd1; Th1: Cxcr6, Tbx21, Slamf1, Entpd1, Gzmb). The Effector cluster expressed Ikzf2, Foxp3, and Ctla4 and was merged with the Treg cluster, giving eight final states. For TCR analysis, productive contigs from Cell Ranger VDJ were imported into R and processed with scRepertoire (Borcherding et al., 2020) together with manual parsing, retaining only productive α/β chains matched to gene-expression barcodes. OT-II cells were identified by TRAV14 (Vα2), TRBV12-1/TRBV12-2 (Vβ5) and CDR3 sequences (CDR3α: CAASRGTGNTGKLIF; CDR3β: CASSSPGQQDTQYF). Cluster marker and between-cluster differential expression was performed with FindMarkers (Wilcoxon; FDR $< 0.05$, $|\log2FC| > 0.25$). Single-cell RNA-seq, TCR-seq, and cell-hashing data have been deposited in GEO# GSE337195.

#### Cross-dataset validation of CD4 T cell subsets

To validate the Stem-like, Tfhl, and Th1 clusters, each cluster was scored using published Tfh, Th1/effector, and progenitor signatures from independent studies. Tfh and Th1 signatures were obtained from: the ProjecTILs CD4 reference atlas of LCMV GP66-specific T cells (Carmona et al., eLife 2022; doi:10.6084/m9.figshare.16592693), chronic LCMV CD4 T cells (Xia et al., Immunity 2022; GSE181474), the TRAMP-C1 prostate model (Cardenas et al., Nature 2024; GSE274801), and nine human cancer types from the Zheng et al. pan-cancer T cell atlas (Science 2021): non-melanoma skin cancer (BCC and SCC; Yost et al., Nat Med 2019), CRC (Zhang et al., Nature 2018), HCC (Zhang et al., Cell 2019), melanoma (Li et al., Cell 2019), NPC (Liu et al., Nat Commun 2021), PDAC (Peng et al., Cell Res 2019), AML (Van Galen et al., Cell 2019), HNSCC (Puram et al., Cell 2017), and lung cancer (Zilionis et al., Immunity 2019). Stem-like/progenitor signatures were derived from the Carmona Tcmp, Xia Tprog, and Cardenas Stem-like populations. For the Carmona reference and Zheng atlas, cluster marker lists were extracted from published supplementary data. For Xia and Cardenas, cluster-specific markers were derived from deposited scRNA-seq data using Seurat FindMarkers (one-vs-two comparisons between Tfh, Th1/Teff, and Stem-like/Tprog; positive markers only; min.pct = 0.25; logfc.threshold = 0.5). Ribosomal protein genes were excluded from progenitor signatures. External signatures were matched to the mOVA dataset by uppercase gene symbol (direct symbol matching rather than formal orthology mapping). The top 30 marker genes per signature were intersected with genes present in the mOVA sorted CD4 dataset, and Seurat AddModuleScore computed per-cell enrichment scores. Mean scores were calculated per cluster (Stem-like, Tfhl, Th1), row z-scored, and visualized as a heatmap (pheatmap) with rows grouped into Tfh, Th1, and Stem-like blocks.

#### Multidimensional scaling (MDS)

To position the mOVA CD4 subsets within a broader landscape of CD4 differentiation across published datasets, MDS was performed on pseudobulk expression profiles. Pseudobulk profiles were generated by summing raw counts across all cells within each cluster (Seurat AggregateExpression), followed by counts-per-10,000 normalization and log1p transformation. The analysis included 57 populations: three mOVA clusters (Stem-like, Tfhl, Th1); nine Carmona LCMV populations (including Th1-Effector, Tfh-Effector, Tcmp, Th1-Memory, Tfh-Memory); three Xia LCMV populations (Tprog, Tfh, Teff); three Cardenas prostate populations (Stem-like, Tfh, Teff); 36 Zheng pan-cancer CD4 populations (Tfh, TfhTh1, Tem, and Temra clusters across nine cancer types); and additional reference populations (NHP germinal-center Tfh, influenza GC Tfh, and CD4-sorted LCMV GP66-specific profiles). Profiles were combined using the intersection of expressed genes and matched by uppercase gene symbol; expression values were then row z-scored within each dataset. A Spearman correlation matrix was computed between all profiles, converted to a distance matrix (1 − correlation), and classical MDS (cmdscale, k = 2) was applied. In place of an explicit batch-correction step, this per-dataset z-scoring minimized species- and platform-specific differences in expression scale, allowing mouse and human populations to group by biological identity rather than study of origin. Points were colored by functional category (Tfh, Th1/Effector, human CXCL13+ TfhTh1, Stem-like, Bulk GP66), with mOVA clusters highlighted as enlarged diamonds.

#### Single-cell TCR and transcriptome analysis of clonal fate

PD-1$^{+}$CD44$^{+}$ activated CD4 T cells were sorted from tumors of Helped (OT-II) and Helped + αPD-L1 (OT-II + αPD-L1) mice and processed for paired single-cell 5′ gene expression and V(D)J (TCR) sequencing (10x Genomics). Transcriptomes were integrated and annotated as described above (Seurat v5, SCTransform integration), yielding the CD4 fate annotations used here (Stem-like, Tfhl, Th1). TCR contigs were assembled with Cell Ranger, and clonotypes were assigned from filtered_contig_annotations.csv using scRepertoire (Borcherding et al., 2020) together with manual parsing in R, retaining only productive chains and matching cell barcodes to the gene-expression metadata. Clonotypes were defined by the endogenous TCRβ CDR3 amino acid sequence (the OT-II transgene was excluded from clone definition), and each cell was assigned the fate of its transcriptional cluster.

#### Clonal-fate heatmap

Endogenous TCRβ clonotypes containing at least one Stem-like cell were retained, and for each clonotype the distribution of its cells across the Stem-like, Tfhl, and Th1 fates was computed (fraction of the clone's cells in each fate, row-normalized per clonotype). Clonotypes were stratified into two condition blocks (Helped and Helped + αPD-L1), labeled by CDR3β sequence and clone size (n), and ordered by clone size within each block. The heatmap was rendered with pheatmap (no clustering) using a diverging blue–white–red scale. The shared clonotype CASSLDRGQDTQYF (n = 15 in Helped, n = 31 in Helped + αPD-L1) is highlighted in both blocks.

#### Shared-clonotype fate composition

The clonotype CASSLDRGQDTQYF, recovered in all six mice (three Helped, three Helped + αPD-L1), was analyzed per mouse. For each mouse, the cells of this clone (Helped: n = 5, 7, 3; Helped + αPD-L1: n = 12, 10, 9) were tabulated by fate and plotted as the percentage of the clone in each of the Stem-like, Tfhl, and Th1 states. The association between treatment and Th1 emergence was tested at the mouse level (Th1 detected in 3 of 3 αPD-L1 mice versus 0 of 3 Helped mice; $P = 0.05$, one-sided Fisher's exact test.

#### Supplementary Figure 1

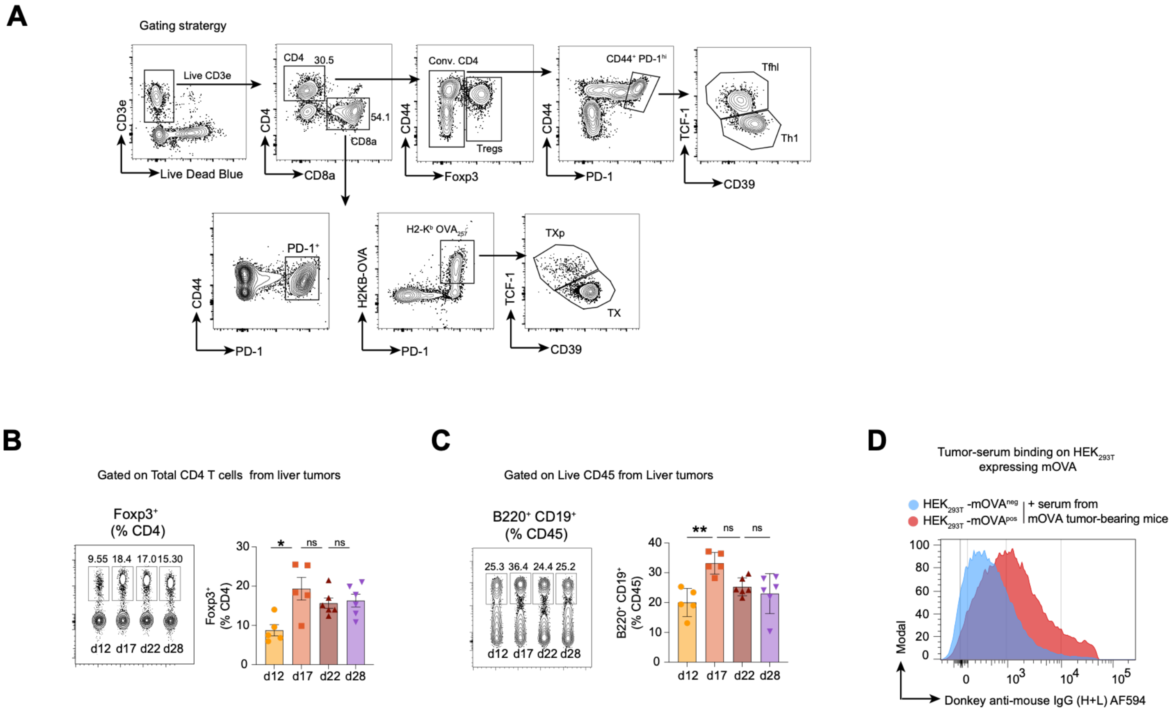


**Fig. S1. Murine model of liver cancer with tractable CD8, CD4 and B cell responses** **(A)** Representative flow cytometry gating strategy used to identify liver-infiltrating CD4 and CD8 T cell subsets. **(B)** Representative flow cytometry plots and graph show Foxp3$^{+}$ cells among liver-infiltrating CD4 T cells at the indicated time points after HDTVI.**(C)** Representative flow cytometry plots and graph show B220$^{+}$CD19$^{+}$ B cells among liver-infiltrating live CD45$^{+}$ cells at the indicated time points after HDTVI. **(D)** Representative histogram shows binding of serum from mOVA tumor–bearing mice (22 days post HDTVI) to HEK293T cells transiently transfected for expression of mOVA, compared with mOVA-negative control HEK293T cells. Symbols represent individual mice. Data are shown as mean ± SEM. Statistical significance was determined by one-way ANOVA with Tukey's multiple comparisons test. **P < 0.01, *P < 0.05. ns = not significant.

#### Supplementary Figure 2

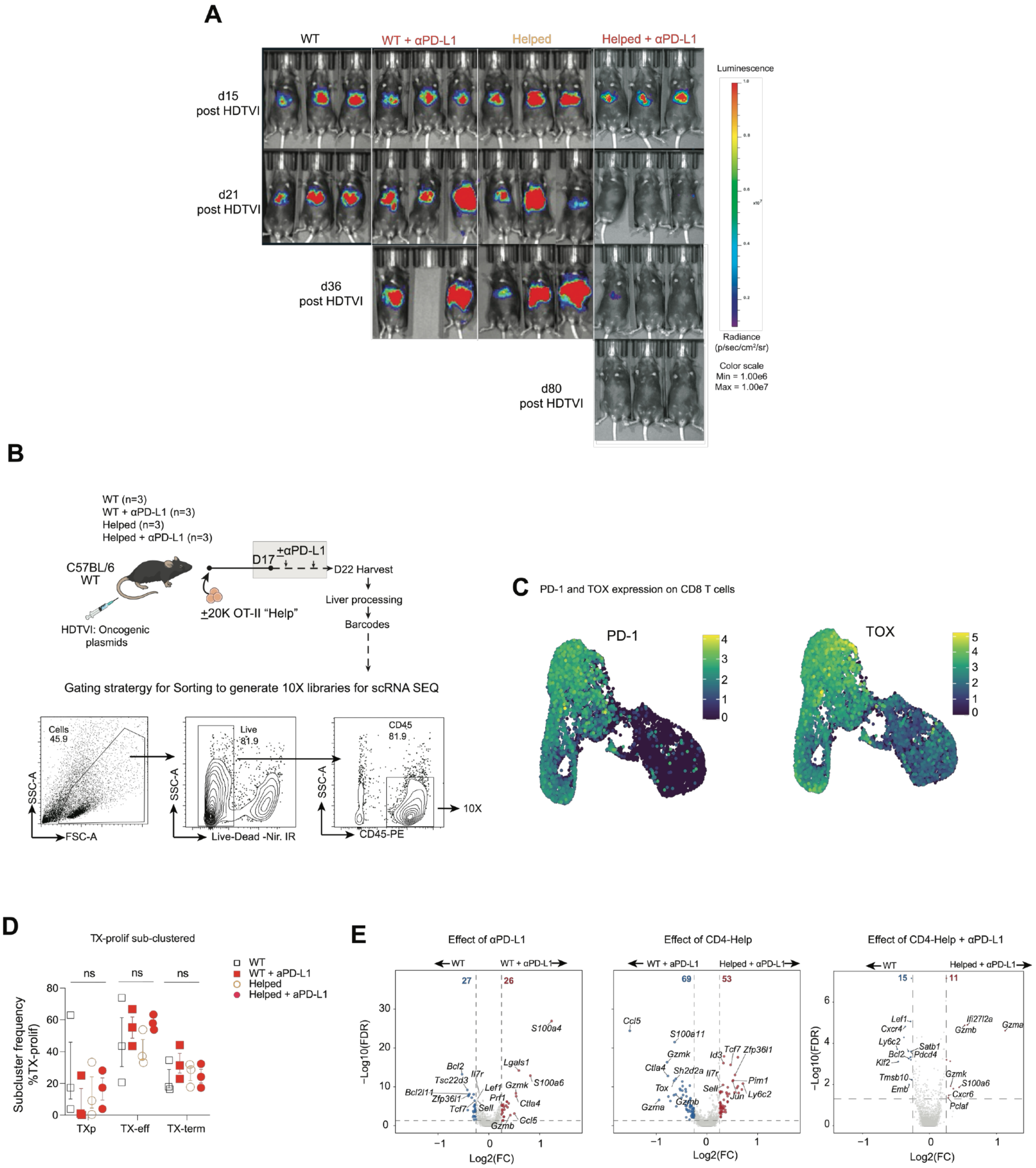

**Fig. S2. CD4 T cell-help sensitizes tumors to overcome resistance to PD-1 blockade**.**(A)** Representative images obtained by In Vivo Imaging System (IVIS) of luciferase activity at the indicated days after HDTVI. Missing animals indicate euthanasia.**(B)** Experimental design: mice received HDTVI on day 0 and 20,000 OT-II-CD4 T cells on day 3, followed by αPD-L1 therapy starting on day 17. Livers were collected at day 22 and processed for single-cell RNA sequencing (scRNAseq). Flow plots show gating strategy for sorting live $CD45^{+}$ cells. **(C)** UMAP feature plots showing PD-1 and TOX expression across CD8 T cells. **(D)** Frequencies of sub-clusters within the TX-prolif CD8 T cell compartment, including TXp, TX-eff, and TX-term states, across the indicated treatment groups. N=3 per group. One-way ANOVA. ns = not significant. **(E)** Volcano plots showing differentially expressed genes (DEGs) in the TX-proliferating cluster to assess effect of αPD-L1, CD4-help during αPD-L1 treatment, and the combined effect of CD4-help plus αPD-L1 compared with untreated mice. Numbers indicate DEGs at False Discovery Rate (FDR) < 0.05 and |log2FC| > 0.25. Red and blue denote upregulated and downregulated, respectively. Select genes are labeled.

### Supplementary Figure 3

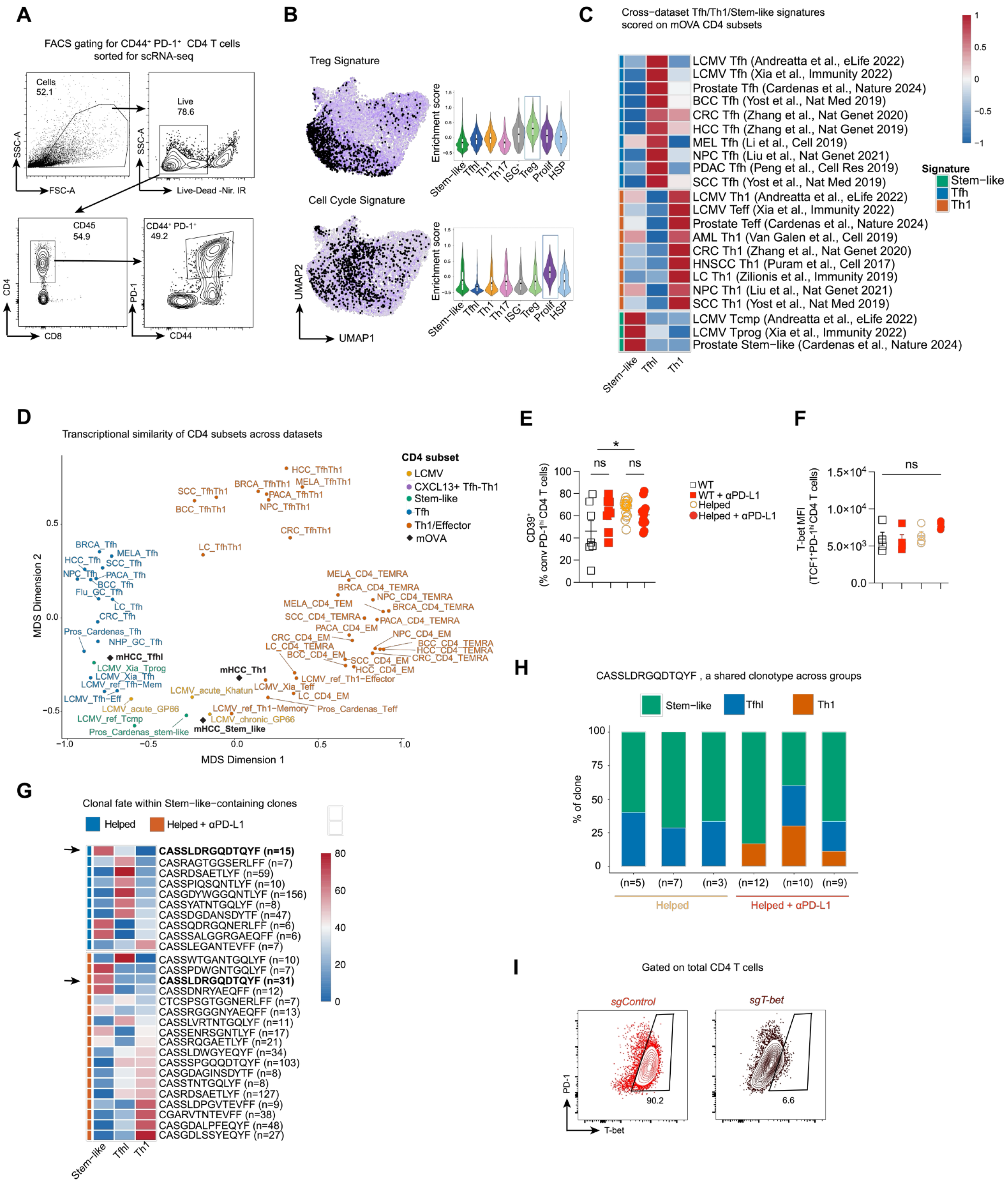

**Fig. S3. Impact of PD-1 blockade on tumor-infiltrating PD-1$^+$ CD4 T cells. (A)** Representative gating strategy for sorting CD44$^+$PD-1$^+$ CD4 T cells from CD4-enriched liver-infiltrating cells for single-cell RNA sequencing. Mice received HDTVI on day 0 and 20,000 OT-II- CD4 T cells on day 3, followed by αPD-L1 therapy starting on day 17. Livers were collected at day 22 and processed for single-cell RNA sequencing (scRNAseq).**(B)** UMAP and enrichment scores showing Treg (top) and cell-cycle (bottom) gene signatures across CD4 T cell clusters. **(C)** Heatmap of published Tfh, Th1, and stem-like CD4 T cell gene signatures scored onto mOVA HCC CD4 subsets (columns: Tfhl, Th1, stem-like clusters). Rows represent signatures derived from LCMV infected mice (*45, 62*), murine prostate cancer model (*25*); and human cancers (signatures from the pan-cancer T cell atlas (*63*)): basal cell carcinoma (*64*), colorectal cancer (*65*), head and neck squamous cell carcinoma (*66*), hepatocellular carcinoma (*67*), lung cancer (*68*), melanoma (*69*), nasopharyngeal carcinoma (*70*), pancreatic adenocarcinoma (*71*), squamous cell carcinoma (*64*), and acute myeloid leukemia (*72*). Signature scores are row-scaled.

**(D)** Multidimensional scaling (MDS) plot of transcriptional similarity between mOVA-tumors-infiltrating PD-1$^+$ CD4 T cell clusters and published CD4 T cell populations across LCMV infection and human cancer datasets. Distance matrix computed as 1 − Spearman correlation of Pseudobulk expression profiles. Each point represents a CD4 subset from a given study, colored by annotated identity. **(E)** Frequency of CD39$^+$ Th1 cells among liver-infiltrating PD-1$^+$TOX$^+$ conventional CD4$^+$ T cells. One-way ANOVA with Tukey's multiple comparisons test. $P < 0.05$. ns = not significant. **(F)** T-bet mean fluorescence intensity (MFI) among TCF1$^+$ PD1$^{hi}$ liver-infiltrating CD4 T cells. One-way ANOVA with Tukey's multiple comparisons. ns = not significant.**(G)** Clonal fate composition of stem-like–containing TCR clonotypes. Each row represents an individual endogenous CD4 T cell clonotype that contains at least one cell in the stem-like cluster. Stacked bars show the proportion of cells assigned to stem-like, Tfhl, and Th1 clusters within each clonotype, stratified by treatment condition (Helped versus Helped + αPD-L1). Clone size (n) is indicated in parentheses.**(H)** Phenotypic fate composition of a shared clonotype CASSLDRGQDTQYF (present across all six mice) shown per individual mouse in Helped (n = 3) versus Helped + αPD-L1 (n = 3) conditions. Stacked bars represent the percentage of cells assigned to stem-like, Tfhl, and Th1 clusters. Numbers below bars indicate clone size per mouse. **(I)** Representative flow cytometry plots show T-bet expression in CD4 T cells following 4 days of *in vitro* stimulation, after OTII CD4 T cells were electroporated with Cas9 ribonucleoproteins (RNPs) containing single guide (sg) RNA targeting *Tbx21* (sgT-bet) or control F8 gene not expressed in T cells (sgControl).

#### Supplementary Figure 4

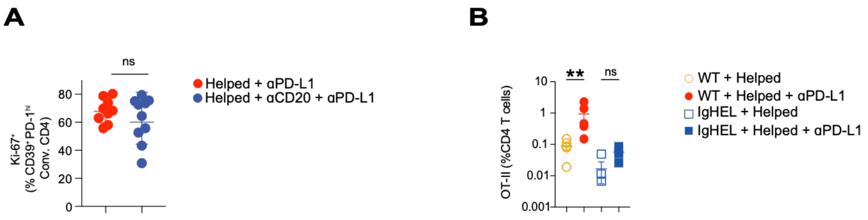


**Fig. S4. Contribution of B cells to effective response to PD-1 blockade (A)** Frequency of Ki-67$^{+}$ among CD39$^{+}$ PD1$^{hi}$ conventional liver-infiltrating CD4 T cells (Th1). C57BL/6 mice received HDTVI on day 0 and 20,000 OT-II- CD4 T cells on day 3, followed by αCD20 (B cells) depleting antibodies on day 15 and 17, and αPD-L1 therapy or isotype control starting on day 17. Flow cytometry analysis was done on day 22. Each point represents an individual mouse. Unpaired Welch's T test. Error bars represent mean ± SEM. ns = not significant. **(B)** Frequency of OT-II cells among liver-infiltrating CD4 T cells. IgHEL transgenic mice or negative littermates (WT) received HDTVI on day 0 and 20,000 OT-II-CD4 T cells on day 3, followed by αPD-L1 therapy starting on day 17. Flow cytometry analysis was done on day 22. Each point represents an individual mouse. ANOVA with Tukey multiple comparisons, Error bars represent mean ± SEM. **$p < 0.01$. ns = not significant.
